# Spatial Regulation of MCAK Promotes Cell Polarization and Focal Adhesion Turnover to Drive Robust Cell Migration

**DOI:** 10.1101/2020.04.30.070557

**Authors:** Hailing Zong, Mark Hazelbaker, Christina Moe, Stephanie C. Ems-McClung, Ke Hu, Claire E. Walczak

## Abstract

The asymmetric distribution of microtubule (MT) dynamics in migrating cells is important for cell polarization, yet the underlying regulatory mechanisms remain underexplored. Here, we addressed this question by studying the role of the MT depolymerase, MCAK, in the highly persistent migration of RPE-1 cells. MCAK knockdown leads to slowed migration and poor directional movement. Fixed and live cell imaging revealed that MCAK knockdown results in excessive membrane ruffling as well as defects in cell polarization and the maintenance of a major protrusive front. Additionally, loss of MCAK increases the lifetime of focal adhesions by decreasing their disassembly rate. These defects are due in part to the loss of the spatial distribution of MCAK activity, wherein activity is higher in the trailing edge of cells compared to the leading edge. Overexpression of Rac1 has a dominant effect over MCAK activity, placing it downstream or in a parallel pathway to MCAK function in migration. Together, our data support a model that places MCAK at a key nexus of a feedback loop, in which polarized distribution of MCAK activity and subsequent differential regulation of MT dynamics contributes to cell polarity and directional migration.

## Introduction

Cell migration is critical for embryonic development, immune response, wound healing, and pathological diseases such as tumor metastasis (Bravo-Cordero et al., 2012; Reig et al., 2014). It is a complex cellular behavior that integrates chemical and mechanical cues from polarized signaling networks, cellular adhesions, and the dynamic actin and microtubule (MT) cytoskeletal networks (Broussard et al., 2008; Kaverina and Straube, 2011; Lauffenburger and Horwitz, 1996; Ridley, 2015). Actin polymerization at the cell front drives the formation of protrusions that provide the force for cell locomotion, and actomyosin exerts contractile forces that move the cell body forward (Cramer et al., 1994). While the contribution of the actin cytoskeleton has been extensively explored, the roles of MTs are relatively understudied.

MTs contribute to multiple aspects of cell migration that are highly coordinated with each other. First, MTs contribute to the directional persistence of migration in several cell types by promoting the initiation and maintenance of cell polarity (Garcin and Straube, 2019; Kaverina and Straube, 2011; Mikhailov and Gundersen, 1998; Takesono et al., 2010; Waterman-Stoer and Salmon, 1997; Zhang et al., 2014). For example, disruption of MT dynamics by nocodazole or taxol slows down cell locomotion by inhibiting the formation of lamellipodia (Liao et al., 1995; Mikhailov and Gundersen, 1998). MT growth also forms a positive feedback loop with signaling molecules, such as the Rho family GTPase Rac1, which promotes actin polymerization and protrusion formation at the leading edge that facilitates cell polarization (Waterman-Storer et al., 1999; Wittmann et al., 2003). Second, MTs spatially and temporarily regulate the assembly and disassembly of integrin-based focal adhesions (FAs) (Broussard et al., 2008; Kaverina et al., 1998; Small et al., 2002; Stehbens and Wittmann, 2012). FAs are transmembrane complexes that couple the actin cytoskeleton to the extracellular matrix, and their dynamic turnover is essential for regulating cellular adhesive properties and intracellular force distribution that impact migration (Ridley et al., 2003). Nascent adhesions form near the protruding leading edge, and most undergo subsequent disassembly in the cell body and trailing edge for cell locomotion (Broussard et al., 2008; Vicente-Manzanares et al., 2009). Targeting MTs to FAs has been shown to promote FA disassembly, possibly mediated by cargo transport to and from the adhesion sites (Broussard et al., 2008; Efimov et al., 2008; Kaverina et al., 1999; Stehbens and Wittmann, 2012). Furthermore, MTs can contribute to migration by interacting with the actin cytoskeleton (Akhshi et al., 2014; Wadsworth, 1999), and by transporting membrane vesicles to the leading edge (Kaverina and Straube, 2011).

Efficient cell movement requires an asymmetric distribution of MT dynamics along the migration axis, which are controlled by numerous MT dynamics regulators in the cell (Etienne-Manneville, 2013; Kaverina and Straube, 2011). Live imaging revealed that stable, long-lived MTs are enriched at the leading edge to support protrusion, whereas more dynamic MTs are found in the cell body and cell rear (Ganguly et al., 2012; Wadsworth, 1999). An example of a spatially regulated MT dynamics modulator in migrating cells is the MT destabilizer, Stathmin/Op18, which was shown to be regionally active in the cell rear (Niethammer et al., 2004). Stathmin is inhibited by phosphorylation via the Rac1-PAK1 cascade, and the level of phosphorylation correlates with the level of cell migration in cancer cells (Liu et al., 2013; Wittmann et al., 2004). Another key MT destabilizer is MCAK (Mitotic Centromere-Associated Kinesin), which was shown to negatively regulate MT polymer levels in interphase (Kline-Smith and Walczak, 2002). In addition, MCAK is overexpressed in numerous cancer and tumor cells, and its expression level correlates with increased lymphatic invasion, metastasis, and poor prognosis (Nakamura et al., 2007; Sanhaji et al., 2011). Previously, MCAK was shown to regulate migration in endothelial cells (Braun et al., 2014), where the authors proposed that a Rac1-Aurora A pathway selectively suppresses MCAK activity at the leading edge to promote regional MT growth (Braun et al., 2014), However, it is not known whether MCAK activity is spatially distributed in migrating cells or what steps of migration are affected by loss of MCAK. Here, we show that MCAK is needed for polarized protrusions to generate a leading edge. MCAK activity is also spatially distributed with higher activity at the trailing edge of cells, consistent with the observations of long stable MTs in the leading edge and more dynamic MTs in the cell body and trailing edge. Overexpression of Rac1 is dominant to loss of MCAK, suggesting that Rac1 acts downstream of MCAK. We postulate a model in which MCAK is a key component of a feedback loop that amplifies the gradient of MT polymerization activity along the migration axis to generate a dominant protrusion during lead edge migration.

## Results

### MCAK is needed for cell migration in RPE-1 cells

To examine MCAK function in cell migration, we tested whether MCAK is required for migration in non-transformed retinal pigment epithelial cells (RPE-1). Using a transwell migration assay, knockdown of MCAK by RNAi resulted in a significant decrease in the relative number of cells that migrated, and a similar reduction in migration was seen with MCAK CRISPR knockout cells (MCAK^-/-^) (Figure 1A-C). As a parallel test, we performed a wound healing assay and found that MCAK knockdown cells were unable to close the wound as robustly as control cells (Figure 1D-E) and had a 17% decreased velocity (Figure 1F). Together, these results indicate MCAK is needed for cell migration in RPE-1 cells. While the MCAK knockout cells would be the ideal system to explore further the effects of MCAK on migration, these cells were phenotypically unstable, as they became large and abnormally shaped after multiple passages. This is likely because depletion of MCAK increases lagging chromosomes that lead to genomic instability (Bakhoum et al., 2009a; Bakhoum et al., 2009b; Kline-Smith et al., 2004). Hence, we pursued the rest of the experiments using RNAi knockdown.

**Figure 1.**
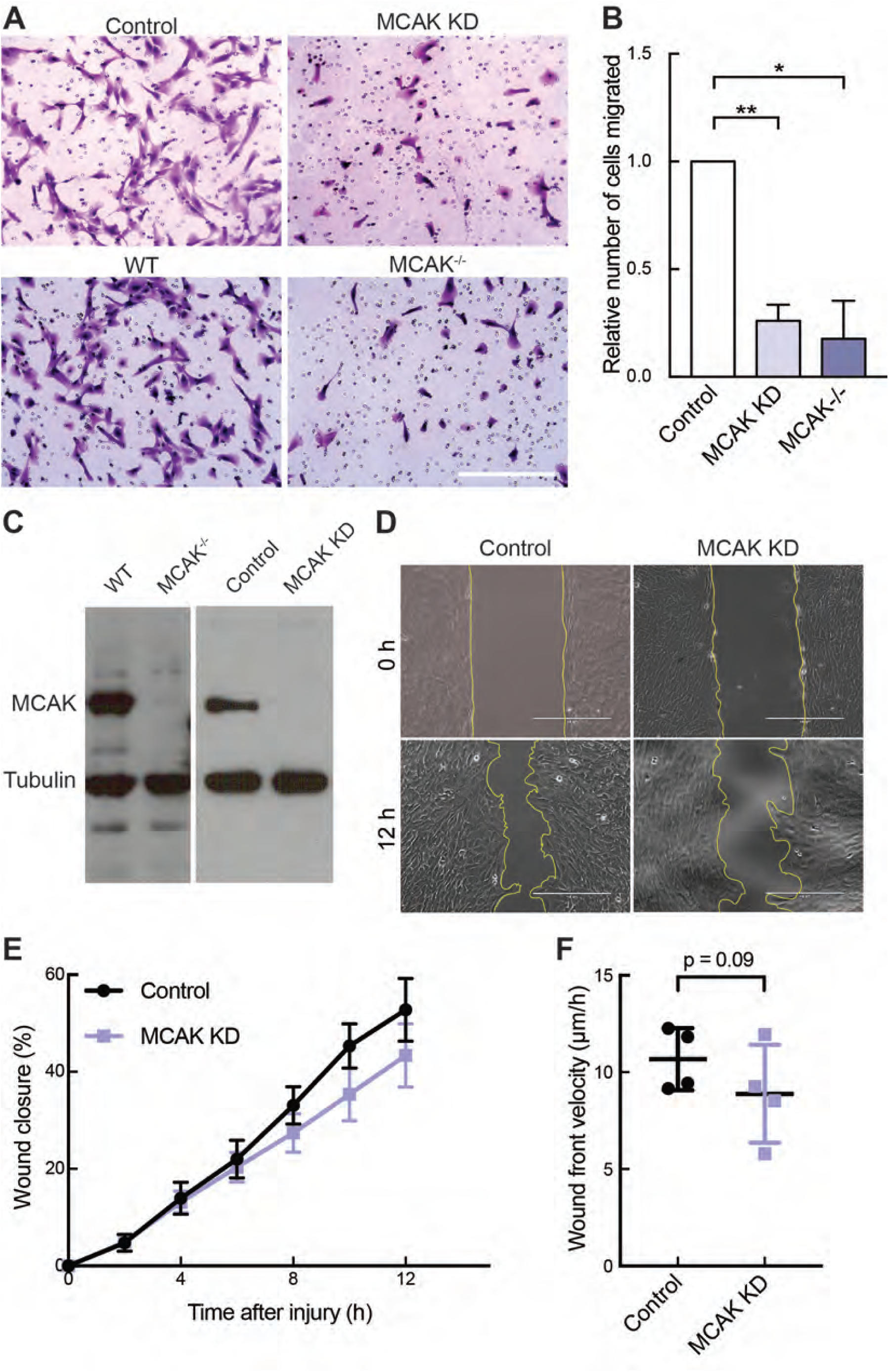
MCAK is required for directional cell migration in RPE-1 cells. **(A)** Representative images of RPE-1 cells stained with Giemsa from the transwell cell migration assay. Scale bar, 200 µm. **(B)** The number of migrated cells per field was counted and normalized to the control (Neg2 siRNA transfection for MCAK knockdown and wild type cells for the MCAK^-/-^ cells). Data represent the mean ± SD from three independent experiments. p values were determined by one-way ANOVA followed by Tukey post-hoc test. *, p<0.05, **, p<0.01. (**C**) Western blot of wild type RPE-1 cells, MCAK^-/-^ RPE-1 cells, or RPE-1 cells treated with control or MCAK siRNAs and probed with a mixture of anti-MCAK and anti-tubulin antibodies. (**D**) Representative phase contrast images of cells at the start and end time points in the wound healing assay. Confluent monolayers of control or MCAK RNAi cells were wounded, and wound healing was imaged every 2 h for 12 h. Wound edges are highlighted by yellow lines. Scale bars, 400 µm. (**E**) Quantification of the percentage of the wound closure by measuring the wound area. Data represents mean ± SEM from four independent experiments. (**F**) Dot plot of the velocity of the advancing wound front determined by linear fitting of wound widths over 12 h. Data represents mean ± SD. p value was determined by two-tailed Student’s *t*-test.

### Loss of MCAK perturbs directional persistence and reduces focal adhesion turnover

To examine how MCAK knockdown impacts the movement of individual cells, we used time-lapse imaging to directly visualize migrating cells on a 2D surface and quantified migration by semi-automatically tracking the nuclei over time. During the 4 h time window examined, most control RPE-1 cells established a polarized morphology with a stable protrusion, and showed highly persistent directional movement typical of RPE-1 cells (Figure 2A top and Figure 2B left) (Zhang et al., 2014). In contrast, ∼30% of cells with MCAK knockdown did not form or maintain a major protrusion but displayed numerous short-lived protrusions with excessive membrane ruffling (Figure 2A bottom and Video 1). These defects in protrusion correlated with reduced migration distance (Figure 2B right) and consequently resulted in a 40% decrease in the overall migration velocity (Figure 2C). It is possible that cell movement was decreased because the individual cells were unable to move effectively and/or because they did not move in a directed fashion. We therefore calculated the instantaneous velocity of the cells during each 10 min imaging interval and found that MCAK knockdown caused a 26% reduction in the median instantaneous velocity (Figure 2D). To address whether the directional persistence of migration was also affected, we calculated the ratio of the displaced migration (Euclidean distance between the start and end points) to the accumulated migration path over time, and found a 24% reduction in the directional persistence (Figure 2E). Overall, these data demonstrate that MCAK is needed to promote efficient movement in RPE-1 cells both by controlling the rate and persistence of movement.

**Figure 2.**
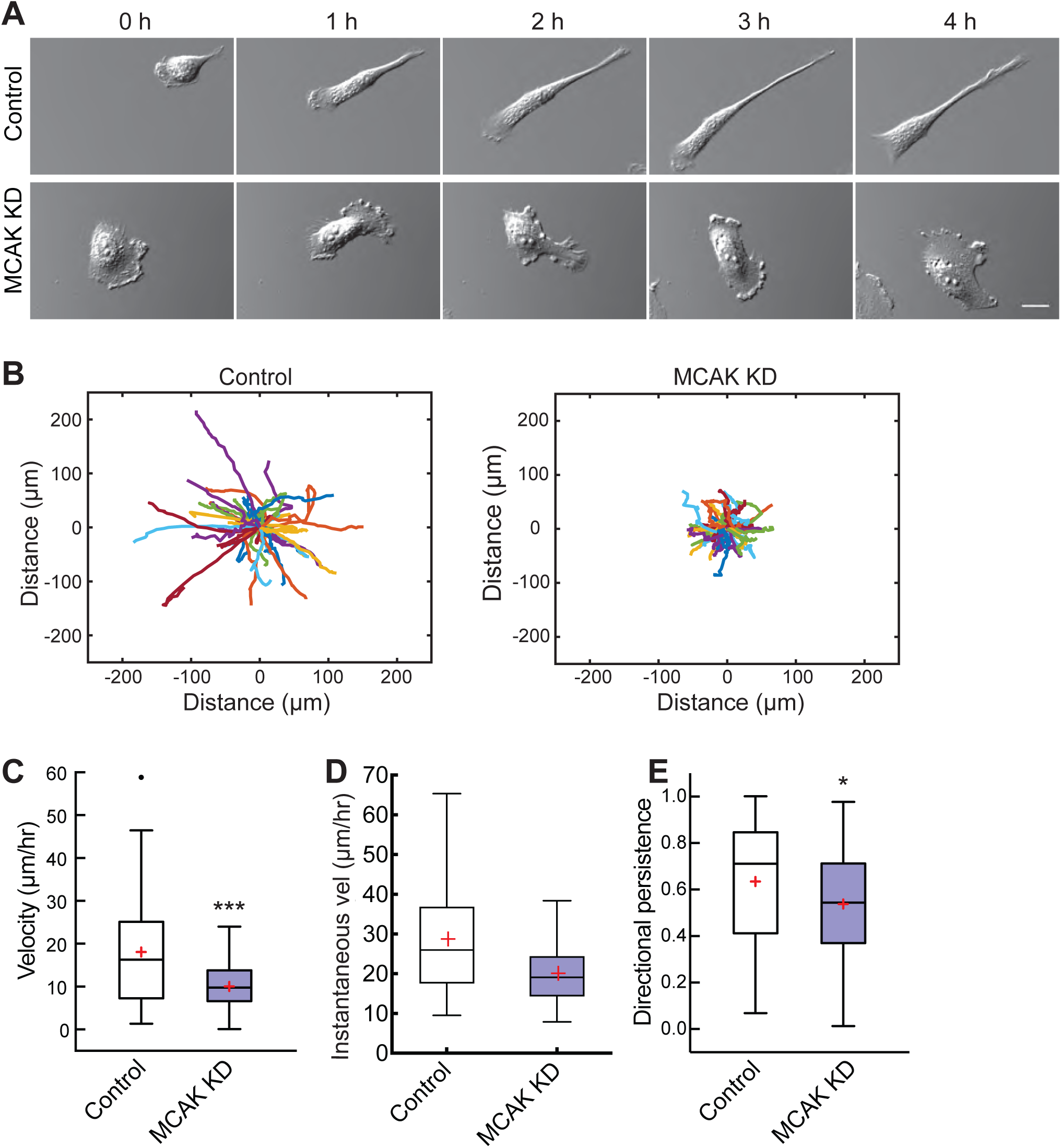
MCAK knockdown results in defective polarized two-dimensional cell migration. **(A)** Representative differential contrast interference images of individual cells from the indicated times during 4 h time-lapse microscopy. Scale bar, 25 µm. **(B)** Pseudo-colored migration tracks of individual cells during the 4 h imaging. Cell position was determined by manually tracking the center of the nucleus over time. n = 60 cells for control, and n = 61 cells for MCAK knockdown. Each colored line represents a single cell. The starting position for each cell was normalized to (0, 0) on the axes. **(C, D)** Quantification of the migration velocity of individual cells determined as the total distance traveled divided by the total time of migration (C) or as the instantaneous velocity of cells during each 10 min imaging interval (D). **(E)** Directional persistence was determined as the displacement divided by the total migration path. Data are represented as a box plot (Tukey), in which the median (line), first and third quartile (box), whiskers (±1.5 times the interquartile range) and mean (red plus sign) are shown. Outliers are indicated as black dots. p values were determined by the two-tailed Mann-Whitney U-test, *, p<0.05, ***, p<0.001.

Because our analysis of cell movement after MCAK knockdown revealed defects in cell morphology, and it is known that MTs are important for the establishment and/or maintenance of cell polarity (Kamath et al., 2014; Schiff and Horwitz, 1980; Takesono et al., 2010), we asked whether MCAK knockdown disrupts cell polarization. Cells were treated with control or MCAK siRNAs for 30 h and then were seeded at low density to allow cells to attach and spread for 20 h before being processed for immunofluorescence. Most control cells showed a classic polarized morphology with a broad front protrusion and a narrow trailing edge (Figure 3A left). In contrast, after MCAK knockdown, many cells displayed a smaller rounded shape without clear polarity (Figure 3A right). In comparison to controls, the median cell area after MCAK knockdown decreased by 43% (Figure 3B), suggesting that cells may have defects in their ability to spread. To compare the cell shape, we measured the eccentricity or roundness of the two populations of cells. Linear objects have an eccentricity of 1, whereas a perfect circular object has an eccentricity of 0. Compared to control cells, MCAK knockdown cells were 20% more round or less eccentric, indicating they were less polarized (Figure 3C). Together, these results show that MCAK contributes to the establishment and/or maintenance of cell polarity, which is a key step for productive migration.

**Figure 3.**
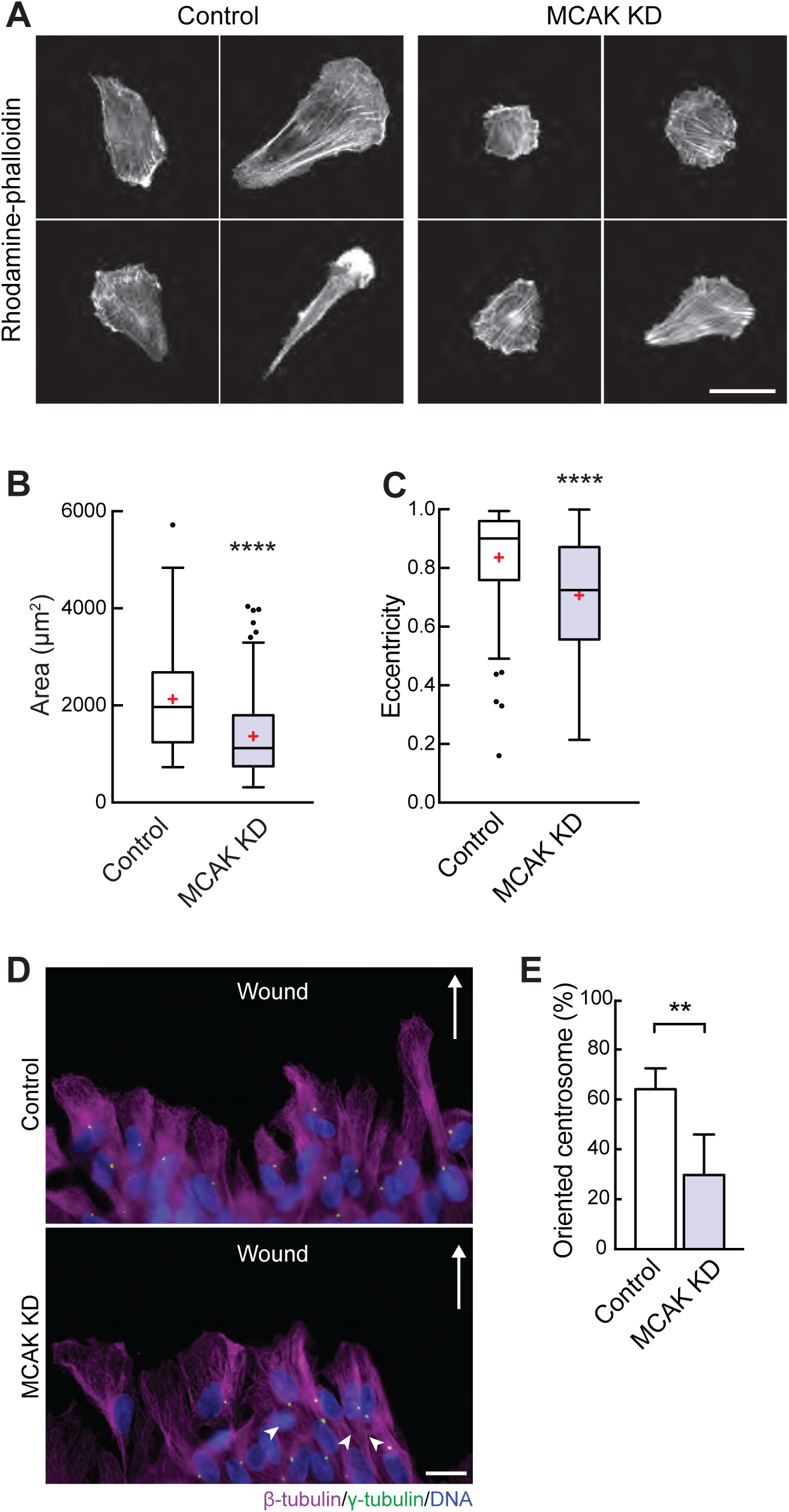
MCAK knockdown causes defects in cell polarization. **(A)** Representative images of individual cells that were grown for 20 h after either control or MCAK knockdown. Cells were stained with Rhodamine-phalloidin to show the actin cytoskeleton. Scale bar, 50 µm. Quantification of the cell area **(B)** and eccentricity **(C)** of cells in (A). n=118 for control and n=150 for MCAK knockdown. Data are represented as a box plot (Tukey), in which the median (line), first and third quartile (box), whiskers (±1.5 times the interquartile range) and mean (red plus sign) are shown. Outliers are indicated as black dots. Data are pooled from three independent experiments. Mann-Whitney U-test, ****, p<0.0001. **(D)** Representative control or MCAK knockdown images of cells at a wound edge. A confluent monolayer of cells was wounded, and cells allowed to migrate for 2 h. Cells were then fixed and stained to visualize MTs (magenta), centrosomes (green), and DNA (blue). The arrows point in the direction of cell movement toward the wound. Scale bar, 20 µm. **(E)** Quantification of the percentage of cells with oriented centrosomes measured by counting the percentage of cells with the centrosome positioned within a 45°angle in front of the nucleus. Data are the mean ± SEM from four independent experiments, with 90-100 cells counted per experiment. Student *t*-test, **, p<0.01.

A key event in cell polarization is centrosome reorientation in which centrosomes relocate from the center of the cell to the front of the nucleus toward the direction of movement (Luxton and Gundersen, 2011). To test whether MCAK knockdown disrupted centrosome reorientation, we performed a scratch wound assay and quantified the relative position of the centrosome to the nucleus (Figure 3D). Control cells had 64% of their centrosomes oriented to the cell front while only 30% of MCAK knockdown cells had orientated centrosomes (Figure 3E), suggesting that the defects in cell polarity after MCAK knockdown are coupled with defects in centrosome repositioning.

The defects in motility after MCAK knockdown are in part due to the inability of the cells to form stable protrusions in the direction of migration. In addition, translocation of the cell body may be affected if the adherence of the cell to the substrate is also perturbed by loss of MCAK. Because MT dynamics influence the turnover of focal adhesions (FAs) (Stehbens and Wittmann, 2012), and MCAK controls MT dynamics, we postulated that the distribution or turnover of FAs might be affected after MCAK knockdown. We first utilized fixed cell analysis of cells after MCAK knockdown and qualitatively analyzed the distribution of MTs and FA using a scratch-wound assay in cells expressing paxillin-GFP (Figure S1). Two hours after wounding, both control cells and cells with MCAK knockdown had a polarized morphology directed at the wound edge. MCAK knockdown cells had a visible increase in the staining intensity of FAs, suggesting that loss of MCAK increased the stability of FAs. To measure the dynamics of the FAs, we next used time-lapse microscopy of these migrating cells using established methods to determine the rates of assembly and disassembly of FAs at the leading edge of cells (Figure 4A-B, Video 2) (Kenific et al., 2016; Stehbens and Wittmann, 2014; Theisen et al., 2012). Briefly, the assembly phase of the paxillin-GFP fluorescence intensity profile was fit with a logistic function (green), the disassembly phase was fit with a single exponential decay function (red), and the lifetime was determined as the amount of time the fluorescence intensity was above half the maximum derived from the assembly and disassembly (black arrow). In comparison to controls, the median FA assembly rate after MCAK knockdown decreased by 17%, and the median disassembly rate was reduced by 45%, resulting in a 52% increase in FA lifetime (Figure 4C). In addition, the fluorescence intensity profiles of FAs after MCAK knockdown often appeared more complex. In control cells, most of the intensity profiles fit a bell-shape curve (56 out of 60 in control), whereas 34% (20 out of 58 in MCAK knockdown) of the FA intensity profiles in MCAK knockdown showed discontinuous disassembly profiles with more than one local maxima (Figure 4B, middle and right). These data suggest that these FAs may undergo cycles of aborted disassembly attempts, possibly due to the inability to sustain FA disassembly (Meenderink et al., 2010; Miranti and Brugge, 2002). Taken together, our data show that MCAK knockdown disrupts FA dynamics and suggest that MCAK-mediated MT dynamics are required to promote efficient FA turnover at the leading edge of migrating cells.

**Figure 4.**
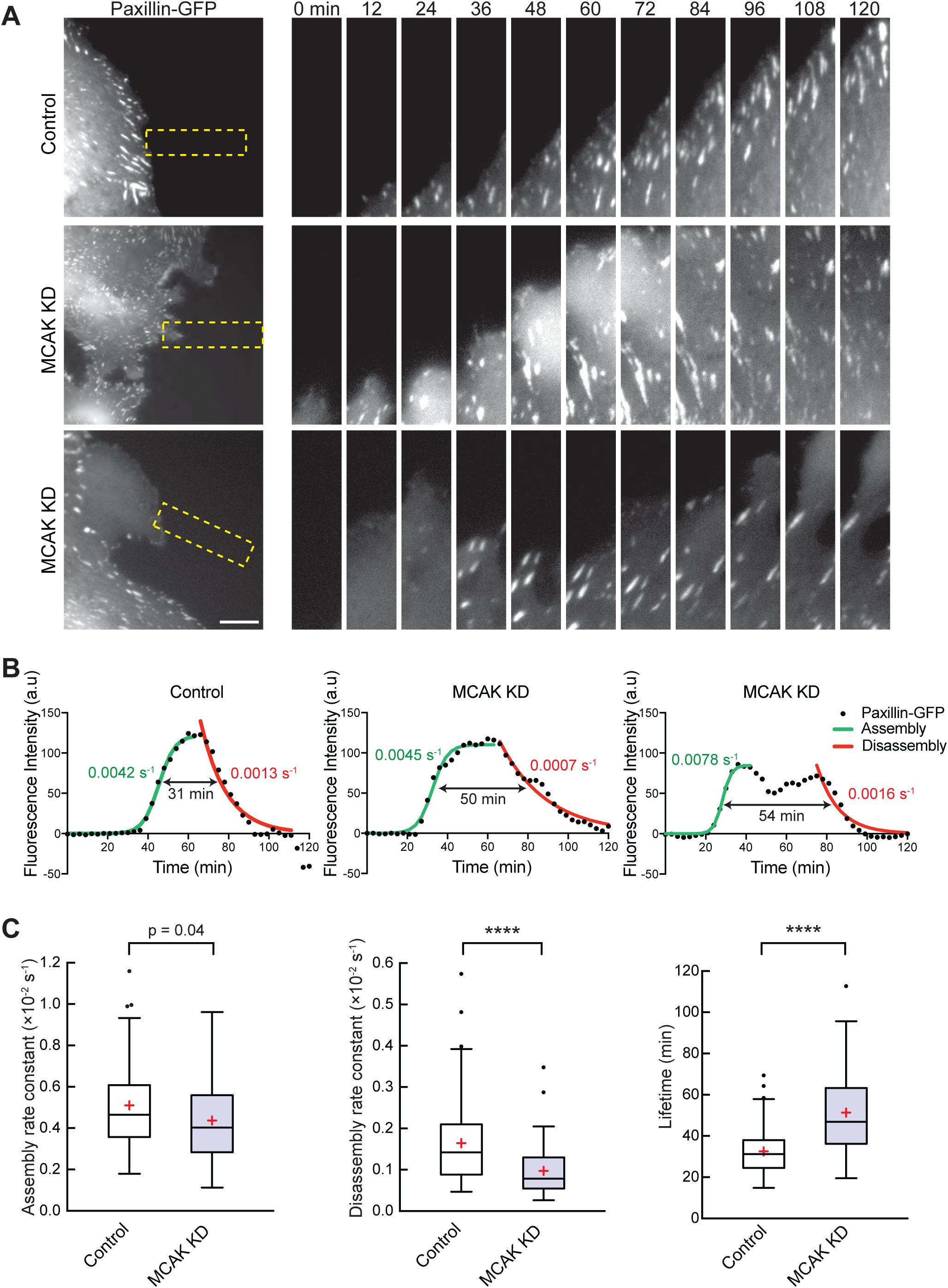
Knockdown of MCAK slows focal adhesion turnover. **(A)** Time-lapse sequences of paxillin-GFP in migrating RPE-1 cells treated with control or MCAK siRNA. Left panels show representative images of cells from control (top) or MCAK knockdown (middle and bottom). Image sequences of the boxed regions are rotated and magnified twofold. Elapsed time in min is indicated above the top panels. Scale bar, 10 μm. **(B)** Representative FA fluorescence intensity profiles of the indicated treatment conditions used for calculating FA turnover parameters in C. Data points are a three-frame running average of FA fluorescence intensity. The green line is a logistic fit for FA assembly, and the red line is an exponential decay fit for FA disassembly. The double arrow indicates the FA lifetime determined by the time in which the fluorescence intensity was above the half-maximum fluorescence intensity of the fit. The assembly rate constant (green), disassembly rate constant (red), and lifetime (black) of each specific profile are shown. **(C)** Quantification of assembly rate constants (left), disassembly rate constants (middle), and lifetime (right) for FAs in cells with control or MCAK knockdown. Data are represented as a box plot (Tukey), in which the median (line), first and third quartile (box), whiskers (±1.5 times the interquartile range) and mean (red plus sign) are shown. n = 66 FAs from 19 cells for control, n = 58 FAs from 20 cells for MCAK knockdown from three independent experiments. p values were determined by the two-tailed Mann-Whitney U-test, ****, p < 0.0001.

### MCAK activity is spatially regulated in cells

In migrating cells, MTs are more stable at the leading edge than in the cell body or trailing edge (Ganguly et al., 2012; Wadsworth, 1999), providing a framework for how cells coordinate the distribution of components by MTs with their actomyosin cytoskeleton for persistent migration. Consistent with this idea, in migrating endothelial cells, it was proposed that MCAK activity is locally inhibited at the cell front by a Rac1-Aurora A pathway to stabilize MT growth (Braun et al., 2014). Aurora A kinase directly phosphorylates MCAK to inhibit its activity (Zhang et al., 2008), and both the conformation and activity of MCAK are reported to be regulated through phosphorylation by Aurora B, which can be assayed by FRET (Förster resonance energy transfer) (Andrews et al., 2004; Ems-McClung et al., 2013; Lan et al., 2004; McHugh et al., 2019; Ohi et al., 2004). To reveal the activity distribution of MCAK activity in migrating cells, we generated a FLIM-FRET biosensor based on its conformational regulation and performed FLIM (Fluorescence Lifetime Imaging Microscopy) on migrating cells in a wound healing assay wherein active MCAK will have a shorter lifetime, and inactive MCAK will have a longer lifetime (Figure 5A). Cells expressing either mEmerald-MCAK donor alone or the FLIM-FRET sensor mEmerald-MCAK-mCherry had mEmerald fluorescence distributed evenly throughout the cells (Figure 5B left), but showed regional differences in lifetimes (Figure 5B right). To compare the differences in lifetime at the leading edge versus the rear of the cell, we drew three 4×4 pixel area boxes at the front of the cell in the direction of the wound edge and at the base of the rear of the cell before the extended tail (Figure 5C). The lifetimes within these regions were averaged and then compared between the control and the sensor. At the leading edge of the cell, the MCAK sensor cells had a similar average lifetime as the control cells (Figure 5D), suggesting that MCAK is in an inhibited state. In contrast, the average lifetime at the rear of the cell was significantly shorter in cells expressing the sensor (Figure 5E), suggesting MCAK is in an active state at the trailing edge. These results are consistent with the idea that MCAK is more active in the tail than at the leading edge, which positions MCAK to play an important role in the regional distribution of MT dynamics for proper cell polarization and persistent migration.

**Figure 5.**
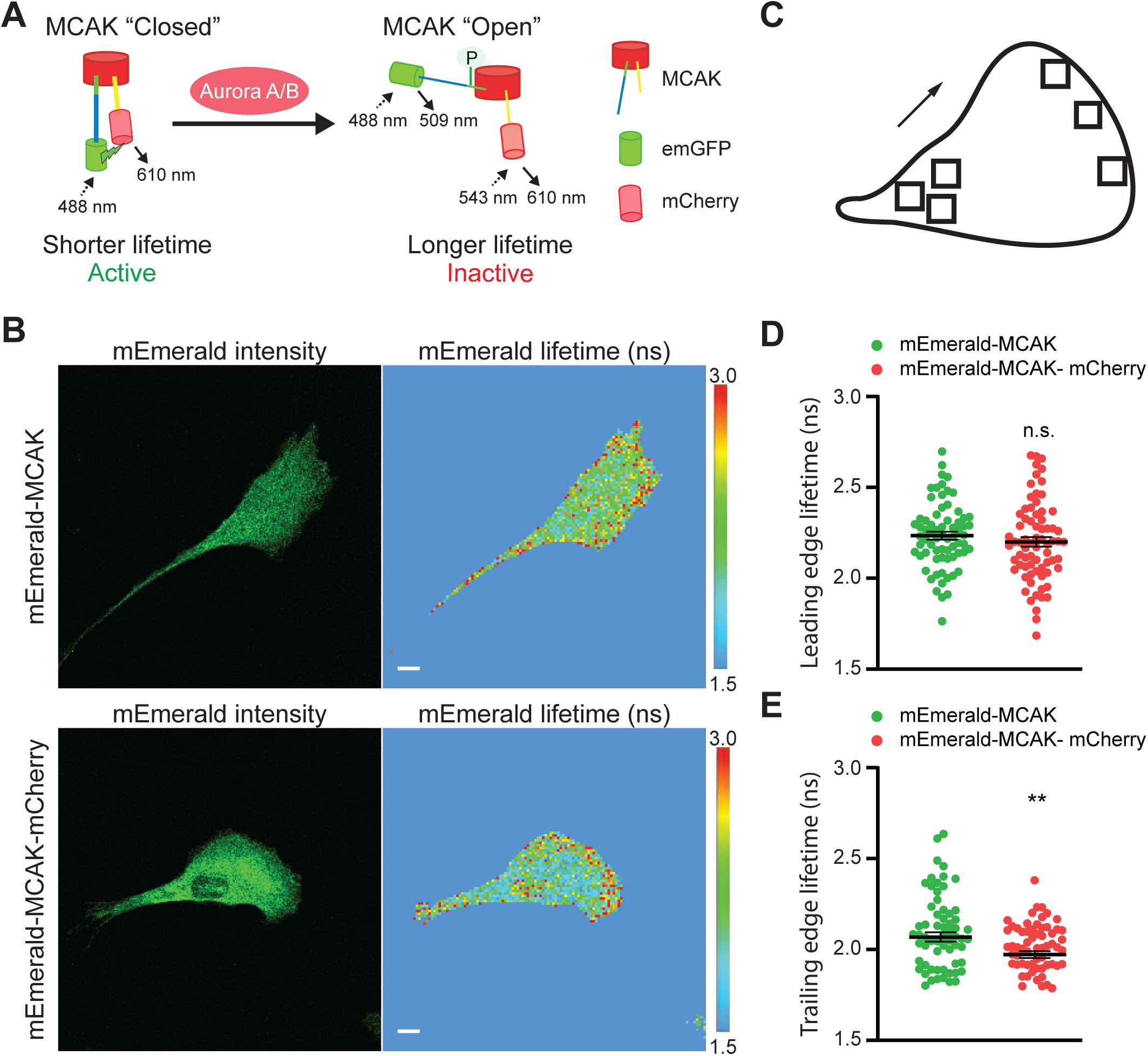
FLIM-FRET reveals that MCAK activity is spatially controlled in migrating cells. **(A)** Schematic illustration of the MCAK-FRET biosensor in which mEmerald is fused to the N-terminus, and mCherry is fused to the C-terminus of MCAK. MCAK in solution is in a closed conformation, which correlates with active MCAK and a short lifetime for the FLIM-FRET sensor. Phosphorylation of MCAK by Aurora A/B changes its conformation from closed to open, which correlates with the inactive MCAK and a longer lifetime for the FLIM-FRET sensor. **(B)** Representative mEmerald fluorescent intensity images (left) and mEmerald fluorescent lifetime measurements (right) of RPE-1 cells expressing the ‘donor-only’ control, mEmerald-MCAK (top), or the FLIM-FRET sensor, mEmerald-MCAK-mCherry (bottom). Pixel-wise fitting of the photon decay rates was done using the tri-exponential tail fit decay model, and the average amplitude weighted lifetime of the mEmerald is shown. The color-coded mEmerald fluorescence lifetime scale is shown at the right. Scale bar, 10 µm. **(C)** Three ROIs were drawn at the leading edge and at the rear of the cell body to calculate the average lifetime in each region. **(D, E)** Dot plots showing the average mEmerald fluorescent lifetimes (ns) found at the (D) leading edge and (E) trailing edge of cells as calculated in (B, C). Lines represent mean ± SEM. n = 69 cells for mEmerald-MCAK and n = 72 cells for mEmerald-MCAK-mCherry from five independent experiments. p values were determined by the Welch’s *t*-test and the two-tailed Student’s *t*-test. N.s., not significant; **, p < 0.01.

### Rac1 acts downstream from MCAK

Cell polarization is coordinated by the activation of Rac1, which promotes actin polymerization, protrusion formation, and membrane ruffling at the leading edge of migrating cells (Hall, 1998; Nobes and Hall, 1999). MT growth stimulates Rac1 activation (Waterman-Storer et al., 1999), which would suggest that increased MT stability from inhibition of MCAK might promote Rac1 activation and cause the MCAK knockdown phenotype we observed in cells. However, studies have suggested that MCAK is downstream of Rac1 in which Rac1 signals through Aurora A dependent inhibition of MCAK (Braun et al., 2014). To ask whether Rac1-induced cell protrusion is enhanced by MCAK knockdown, we transiently expressed constitutively active (CA) GFP-tagged Rac1 in cells after MCAK knockdown and then examined the morphology of cells as described above. Expression of GFP slightly lowered the area of control cells relative to previously observed control cells (Figure 3), which was likely due to differences in the timing of experiments, but GFP expression did not alter the significant difference in eccentricity of control cells versus MCAK knockdown cells (Figure 6A top and 6B-C). In contrast, expression of CA-GFP-Rac1 enhanced cell spreading as evidenced by broad lamellipodia in both control and MCAK knockdown cells (Figure 6A bottom). Quantification of the morphological parameters confirmed that CA-GFP-Rac1 expressing cells showed dominant effects of increased cell area and reduced eccentricity, in both control and MCAK knockdown cells (Figure 6B-C). These results suggest that Rac1 activation occurs downstream of MCAK or in a parallel pathway, and that the excess transient protrusions seen after MCAK knockdown may result from local Rac1 activation by enhanced MT growth that occurs in the absence of MCAK. These results also raise the possibility that MCAK is a key component of a feedback loop that mediates MT growth at the leading edge of migrating cells.

**Figure 6.**
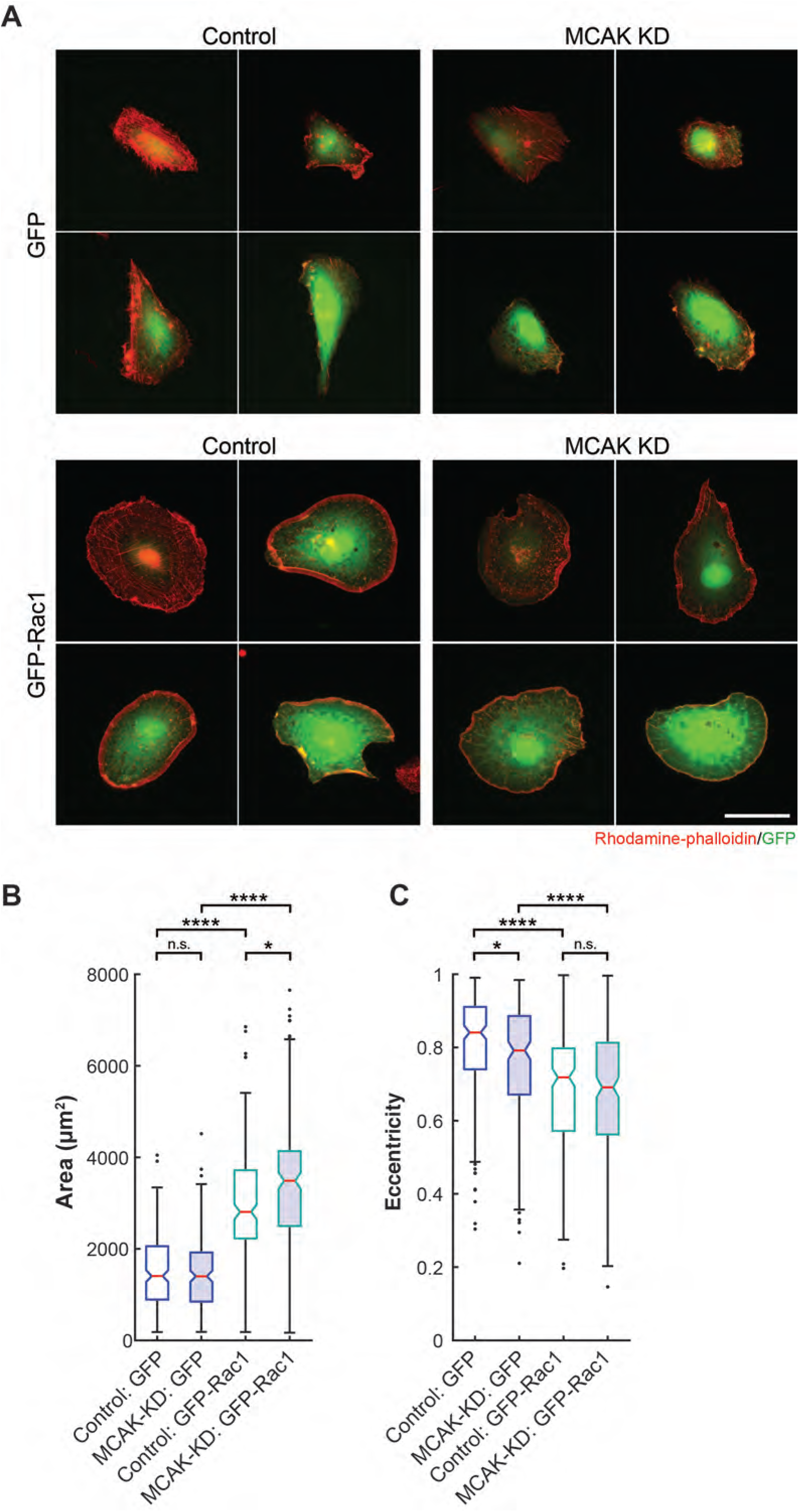
Rac1 acts downstream of MCAK. **(A)** Representative images of RPE-1 cells with control or MCAK knockdown in which cells were plated at low density on coverslips and then allowed to adhere and spread for 20 h. Cells were then transiently transfected with a plasmid expressing GFP or CA-GFP-Rac1 (GFP-Rac1). At 6 h post-transfection, cells were stained with Rhodamine-phalloidin to show the actin cytoskeleton. Images are scaled equivalently. Scale bar, 50 µm. Quantification of the cell area **(B)** and eccentricity **(C)** of cells in (A). Data are pooled from five independent experiments. n = 201 for Control: GFP, n=185 for MCAK-KD: GFP, n = 219 for Control: GFP-Rac1, and n=206 for MCAK-KD: GFP-Rac1. Data are represented as a box plot (Tukey), in which the median (line), first and third quartile (box), whiskers (±1.5 times the interquartile range) and mean (red plus sign) are shown. p values were calculated by the Kruskall-Wallis test followed by Dunn’s post-hoc test for multiple comparisons. *, p<0.05, ****, p<0.0001, n.s., not significant.

## Discussion

In this study, we showed how the spatial regulation of MT dynamics by MCAK contributes to directional cell migration (Figure 7A). Abolishing MCAK activity disrupted cell migration by negatively affecting several important aspects, including cell polarization/protrusion formation, centrosome reorientation, and the turnover of focal adhesions (Figure 7B). Because MCAK needs to be regionally inhibited at the leading edge of a migrating cell to promote MT growth (Braun et al., 2014), our results highlight the importance of the tight spatial control of MCAK activity to generate and maintain one dominant protrusion for directional movement.

**Figure 7.**
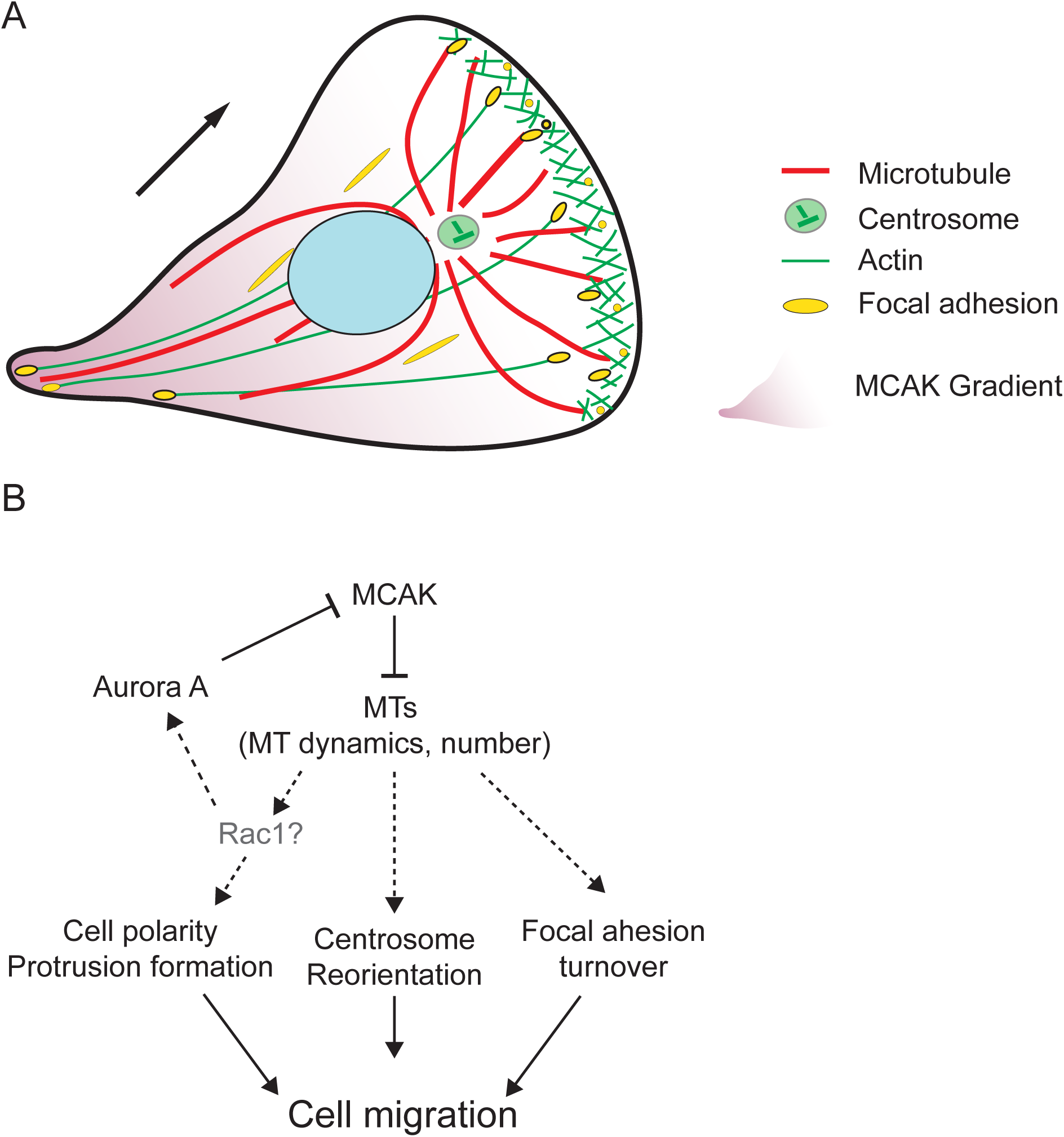
MCAK-regulated MT dynamics affects directional cell migration. **(A)** Cells show highly persistent cell migration on 2D surfaces with a dominant protrusion in the cell front that is dependent on MTs, actin, and focal adhesions. The work shown here suggests that spatial control of MCAK activity is needed for cell polarization and persistent migration. **(B)** MCAK down regulation is known to disrupt MT dynamics and increase the number of MTs, which may result in retarded cell migration due to combined defects in cell polarization, extra protrusion formation, centrosome reorientation, and slower FA turnover.

One emerging idea in the field of cell migration is that cytoskeletal dynamics need to be spatially controlled, but the molecular mechanisms demonstrating differences in local concentrations of protein or a spatial distribution of their activity have not been well-established. Our work takes advantage of an MCAK FRET-based biosensor, which reports conformational changes in MCAK that correlate with its activity (Ems-McClung et al., 2013). Thus, the present findings represent the first demonstration of spatial changes in MCAK activity at the leading and trailing edges of cells, which are likely critical for localized differences in MT dynamics for polarized cell migration.

Our results evoke new thoughts about how localized MT dynamics are coupled to the formation of a stable leading edge. Loss of MCAK results in enhanced membrane ruffling and short-lived protrusions, which are likely due to the stabilization of MTs due to the lack of MCAK-induced MT depolymerization. Consistent with this idea, stabilizing MTs by taxol treatment leads to the formation of membrane protrusions evenly distributed around the cell perimeter rather than mainly at the cell front (Schiff and Horwitz, 1980; Waterman-Storer and Salmon, 1999). Down-regulation of the MT destabilizer, Stathmin/Op18, also enhances membrane ruffles in macrophage activation (Xu and Harrison, 2015), supporting the idea that global stabilization of MTs disrupts polarity. The enhanced formation of random protrusions and membrane ruffling upon the depletion of MCAK indicates that MCAK might be involved in regional suppression of signals to prevent the formation of extra protrusions around the periphery, thus ensuring a dominant lamellipodial protrusion for efficient migration. MT growth locally activates Rac1, which in turn promotes protrusion formation (Waterman-Storer et al., 1999); thus it is possible that MCAK knockdown results in enhanced global MT growth, which promotes Rac1 localization leading to short lived protrusions and ruffling all over the cell cortex. In support of this idea, our data showed that most CA-Rac1-GFP transfected cells have broad lamellipodia in both control and MCAK knockdown cells, suggesting that Rac1 acts downstream of MCAK, although it is also possible that Rac1 acts in a parallel pathway. A previous study postulated that Rac1 was upstream of MCAK and worked through a Rac1-Aurora A pathway to locally inhibit MCAK MT depolymerization activity (Braun et al., 2014). One model that could reconcile both observations is that there is a feedback loop in which Rac1 stimulates protrusion formation, which signals to further locally inhibit MCAK, resulting in MT stabilization and persistent cell polarization (Figure 7B). These stabilized MTs then may serve as tracks on which material is transported to the leading edge of cells to generate a single dominant lamellar protrusion.

It is currently not clear whether the defects in centrosome position are a direct or indirect result of MCAK inhibition. In some cell types, movement of the centrosome during migration is correlated with the establishment of the leading edge. Because inhibition of MCAK enhances multiple protrusions and disrupts directional migration, it is possible that the failure to establish a single dominant protrusion consequently results in the failure to properly position the centrosome. Alternatively, it has been shown that overexpression of MCAK activates the release of MTs from the centrosome (Ganguly et al., 2011a; Ganguly et al., 2011b), so perhaps inhibition of MCAK suppresses release of MTs from the centrosome, which changes the organization of MTs in that region and prevents centrosome repositioning. In support of this idea, we also noted in earlier work that there was an accumulation of MTs in the perinuclear area after MCAK inhibition (Kline-Smith and Walczak, 2002), suggesting that MCAK may act on short MT plus ends near the centrosome, or that MCAK activity may also be important at MT minus ends.

Another key component in migration is the formation and release of FAs, which provide traction for motility. We showed that MCAK knockdown increased leading-edge FA lifetime and disturbed FA turnover mainly by reducing the disassembly rate resulting in more persistent FAs. It was shown previously that MT targeting and subsequent catastrophe facilitate FA disassembly (Efimov and Kaverina, 2009; Efimov et al., 2008). MTs may provide tracks for the transport of cargos that are involved in FA assembly and disassembly to and from the adhesion sites (Seetharaman and Etienne-Manneville, 2019; Stehbens and Wittmann, 2012). For instance, integrin-linked kinase is required for MT stabilization and caveolin transport to FAs (Wickstrom et al., 2010). Caveolin mediates integrin endocytosis that may facilitate FA disassembly (Kiss, 2012; Stehbens and Wittmann, 2012). In addition, the MT-dependent transport and accumulation of matrix metalloproteases at FAs promote exocytosis that degrades ECM and consequently facilitates FA disassembly (Takino et al., 2006; Wiesner et al., 2010). Because MCAK does not associate directly with FAs but rather with the plus-ends of MTs ((Moore et al., 2005) and our unpublished results), where it controls their dynamics (Braun et al., 2014; Kline-Smith and Walczak, 2002), we favor the idea that the increased MT stability generated by the loss of MCAK indirectly affects the transport of these other components along the MTs that interact with FAs. If we assume that MCAK knockdown does not change the level of other proteins in cells, an overall increase in the number of MTs may result in fewer cargos per MT, which could lead to lower efficiency in disassembly of FAs. This “dilution effect” may also account for our observed the slight decrease in the assembly rate.

Previously, it was reported that several MT plus tip tracking proteins, APC, CLASPs, and ACF7, in addition to their activity in MT remodeling, may mediate a physical link between the MT plus ends and FAs, and contribute to robust cell migration (Stehbens et al., 2014; Watanabe et al., 2004; Wen et al., 2004; Wu et al., 2008; Wu et al., 2011). MCAK also tracks MT plus ends (Honnappa et al., 2009; Moore et al., 2005). Plus tip tracking proteins rely on polymerizing MTs to track MT ends; thus the loss of MCAK activity on MT plus tips may cause a change in the composition, dynamics, and/or architecture of the protein complexes riding on growing MT tips due to increased polymerization that results in our observed FA defects.

The defects in FA dynamics after MCAK knockdown may also help explain the defects in cell movement in MCAK RNAi cells, as properly regulated FA dynamics are required for productive forward translocation of the cell body (Stehbens and Wittmann, 2012). Our analysis also showed that the overall decrease in migration after MCAK knockdown was due at least in part from the short-term movement of the cell body. If the cell body is more adherent to the substrate because of slowed FA turnover, this would result in failure of the cell body to be able to readily translocate forward. In addition, we also consistently observed ‘blebbing’ in cells with MCAK knockdown that occurs around the edges of the adhered cell body. It has been proposed that bleb formation is driven by hydrostatic pressure generated in the cytoplasm by the contractile actomyosin cortex, which is myosin-II dependent (Paluch and Raz, 2013). MCAK regulation of MT dynamics has also been shown to be mechanosensitive in a myosin-II-dependent manner in 3D cultured HUVEC cells (D’Angelo et al., 2017), indicating the cross-talk of the two pathways. Thus, it is possible that the ‘blebbing’ phenotype we observed was caused by a change in myosin-II activity after MCAK knockdown. Perhaps this phenotype becomes more apparent in cells that are unable to generate a single leading protrusive edge due to changes in adhesion as well as changes in the contractile actomyosin cortex.

Taken together our results support a model wherein MCAK contributes to overall cell migration at multiple levels (Figure 7). Cells require MCAK to be locally inhibited to allow for the establishment of a single protrusive front with stabilized MTs (Figure 7A). The stabilization of this front is further enhanced by the transport of Rac1 and other factors along the MTs to the leading edge of the cell. MCAK, located at the plus-tips of MTs, acts to ensure the dynamic turnover of these MTs ends, generating a dynamic array of MTs, some of which can interact with FAs and stimulate their turnover, which will in turn allow for the cell body to translocate forward. MCAK may also act on MTs near centrosomes, where it enhances proper centrosome positioning toward the leading edge of the cell. In this model, MCAK control of MT dynamics plays a critical role in a feedback loop that ensures the development of a dominant leading edge for directed cell migration (Figure 7B).

## Materials and Methods

### Cell culture and RNAi

Human retinal pigment epithelial cells immortalized with hTERT (hTERT-RPE-1, ATCC) were cultured in DMEM or Opti-MEM medium (Invitrogen) supplemented with 10% fetal bovine serum, 2 mM L-glutamine, and 50 mg/ml penicillin/streptomycin at 37°C, 5% CO_2_ in a humidified incubator. A stable RPE-1 cell line expressing paxillin-GFP (RPE-1 PG6) was obtained from the laboratory of A. Straube (Theisen et al., 2012). HEK-293 cells were cultured in Gibco(tm) DMEM, High Glucose supplemented with GlutaMAX(tm) and pyruvate (Thermo Fisher), 10% fetal bovine serum (Corning), and 50 mg/ml penicillin/streptomycin at 37°C, 5% CO_2_ in a humidified incubator. Cell line identity was verified by STR profiling.

For siRNA-mediated MCAK knockdown, 3 ×10^5^ RPE-1 cells were seeded in a six-well plate. After 24 h, cells were transfected with 10 nM RNAi oligonucleotides using Lipofectamine RNAiMax (Invitrogen) according to the manufacturer’s instructions. The following siRNAs (Dharmacon) were used: negative control-2 (5’-UGGUUUACAUGUUGUGUGA-3’) and MCAK RNAi (5’-GAUCCAACGCAGUAAUGGU-3’) (Hedrick et al., 2008). Cells were examined at 48-60 h after knockdown depending on the specific type of experiment.

### DNA constructs, transfection, and lentivirus production

Plasmids used for transfection include pEGFP-C1 (GenBank: U55763.1, Clonetech) and pcDNA3-EGFP-Rac1-Q61L (a gift from G. Bokoch, Addgene plasmid #12981) (Subauste et al., 2000). mEmerald-MCAK and mEmerald-MCAK-mCherry were cloned using Gibson assembly by fusing the cDNA of mEmerald, hMCAK and/or mCherry into the backbone of pEGFP-C1 (Clonetech). To generate pLOVE-mEmerald-MCAK and pLOVE-mEmerald-MCAK-mCherry, mEmerald-MCAK and mEmerald-MCAK-mCherry, sequences were cloned into the pENTR/D-TOPO backbone (Invitrogen) and then recombined into the pLOVE plasmid (Addgene #15948) according to the manufacturer’s instructions. For Rac1 overexpression, at 30 h post-RNAi transfection, cells were plated at ∼ 2×10^4^ cells/ml on poly-L-lysine coverslips and allowed to attach for 24 h before DNA constructs were introduced into cells using Lipofectamine 3000 (Invitrogen) according to the manufacturer’s instructions. Cells were analyzed at 6 h post DNA transfection.

Lentiviruses were produced by co-transfecting 1 μmol of pLOVE-mEmerald-MCAK or pLOVE-mEmerald-MCAK-mCherry with the packaging plasmids dRT-pMDLg/pRRE (Addgene #60488), pRSV-Rev (Addgene #12253), and pMD2.G (Addgene #12259) using Lipofectamine 3000 in HEK293T cells plated at 3.5 × 10^6^ cells/ml. Media was changed 6 h post-transfection, and media containing virus was collected at 24 h and 48 h post-transfection, filtered, and stored at −80°C in single use aliquots.

### Generation of MCAK null mutant

An MCAK-/- cell line was generated using CRISPR/Cas9-mediated homology-directed recombination (Ran et al., 2013). CRISPR target sites adjacent to the start codon were selected by CRISPR Design (http://crispr.mit.edu/), and guide RNAs (gRNAs) were designed and cloned into the pSpCas9n(BB)-2A-Puro (PX462) V2.0 plasmid (a gift from Feng Zhang, Addgene plasmid #62987). Homology arms containing the 5’-UTR, mEmerald, and the first exon and intron of MCAK genomic DNA were cloned into the pENTR/D-TOPO plasmid (Invitrogen) by overlap extension PCR. RPE-1 cells were transfected with 1 µg of PX462-gRNA plasmid and 1 µg repair template with Lipofectamine 3000 and were selected with 5 µg/ml puromycin. Single cells were sorted, and positive cell clones were screened by PCR against the inserted genomic region and were further verified by DNA sequencing. The insertion of mEmerald before the first exon of MCAK abolished the gene product of MCAK, as determined by Western blot.

#### gRNA 1

Sense oligo: 5′-AAA CAA TGG CCA TGG ACT CGT CGC C -3′.

Antisense oligo: 5′-CAC CGG CGA CGA GTC CAT GGC CAT T -3′.

#### gRNA 2

Sense oligo: 5′-CAC CGA TCA AGA TCC AAC GCA GTA A-3′;

Antisense oligo: 5′-AAA CTT ACT GCG TTG GAT CTT GAT C-3′.

#### Homology arm 1

F primer: 5’-CAC CGA GAA CGG GCC ATG ATG ACG ATG GCG GTT TT -3’. R primer: 5’-AAA ACC GCC ATC GTC ATC ATG GCC CGT TCT CGG TG -3’.

#### Homology arm 2

F primer: 5’-CTG CCC ATT CCA CTG AAA CAC AGG ATT TCT CCA A -3’ R primer: 5’-TTG GAG AAA TCC TGT GTT TCA GTG GAA TGG GCA G -3’

### Immunofluorescence and Western blotting

For immunofluorescence, cells on coverslips were fixed for 20 min in 4% paraformaldehyde (Thermo Scientific #28908) in phosphate-buffered saline (PBS; 137 mM NaCl, 2.7 mM KCl, 10 mM Na_2_HPO_4_, 1.8 mM KH_2_PO_4_, pH 7.2) and permeabilized for 20 min with 1% Triton X-100 in PBS. Cells were blocked in Abdil-Tx ((TBS; 20 mM Tris, 150 mM NaCl, pH 7.5) with 0.1% Triton X-100, 2% bovine serum albumin, and 0.1% sodium azide) for 30 min at room temperature (RT). Cells were stained with primary antibodies for 1 h at RT or overnight at 4°C, and stained with secondary antibodies and/or phalloidin-rhodamine (1:1000, Santa Cruz #362065) for 1 h at RT. DNA was stained with 10 μg/ml Hoechst for 20 min (Sigma-Aldrich). Coverslips were washed three times with TBS-Tx (TBS with 0.1% Triton X-100) between each step and mounted in ProLong Diamond (Invitrogen). The primary antibodies used were: mouse anti-γ-tubulin (1µg/ml, Sigma-Aldrich GTU-88), mouse DM1α (1:5000, Sigma-Aldrich T6199), or rat anti-tubulin YL1/2 (10 µg/ml, purified in house from the rat monoclonal cell line). Secondary antibodies used were: DyLight488 donkey anti-mouse (1 µg/ml, Jackson ImmunoResearch #715-545-150), DyLight594 goat anti-rat (1.5 µg/ml, Jackson ImmunoResearch #112-515-175), or Alexa Fluor 594 goat anti-mouse (1 µg/ml, Invitrogen #A10239).

For Western blots, cells were collected, washed with PBS, and lysed in 2X sample buffer (0.125 M Tris, 4% SDS, 20% glycerol, 4% β-mercaptoethanol, and a trace amount of bromophenol blue, pH 6.8) at ∼ 1×10^7^ cells/ml. Equal amounts of cell lysates were electrophoresed on 10% (v/v) SDS-PAGE gels and transferred to nitrocellulose (Schleicher & Schuell). Blots were incubated in blocking buffer (TBS with 0.1% Tween 20, 5% (w/v) nonfat dry milk) and probed with mouse DM1α (1:5000), and rabbit anti-hMCAK-154 (0.8 μg/ml) (Hedrick et al., 2008) diluted in Abdil-T (TBS with 0.1% Tween-20, 2% bovine serum albumin, and 0.1% sodium azide). Secondary antibodies were used at 1 μg/ml for goat anti-rabbit and sheep anti-mouse linked horseradish peroxidase (Invitrogen). Blots were developed with SuperSignal West Pico Chemiluminescent Substrate (Pierce).

### Transwell migration, wound healing, and random migration assays

For transwell migration assays, 5×10^4^ cells in serum free DMEM were introduced to the upper chamber of an 8 μm pore size insert (BD Falcon), and DMEM medium supplemented with 10% fetal bovine serum was used as a chemoattractant in the lower chamber. Cells were allowed to migrate for 18 h before they were fixed in 4% paraformaldehyde in PBS for 30 min. Cells were stained with Giemsa (5% diluted in water, Sigma-Aldrich) overnight. The stained inserts were imaged with a 20x objective (0.60 NA, Plan Apo) using an EVOS FL Auto microscope (Life Technologies). The number of cells per image was quantified with a custom MATLAB code. Briefly, 12-bit images were converted to 8-bit grayscale images, local contrast adjusted, and complemented to binary images. Adaptive filtering was used to remove noise, followed by a background cutoff, image fill and open, and a size cutoff of the nucleus. To separate adjacent cells, local maxima that approximately correspond to the cell nuclei were identified, were transformed to local minima and were superimposed over the binary image. All source codes can be found at https://github.com/haizong/cell_morph_tracker. Three triplicate inserts were assayed per condition in an experiment, and the mean of five representative fields per insert was calculated. The results of an individual experiment were averaged from the mean number of cells migrated for the three inserts, and then the assay was repeated on three biological replicate experiments. For the ease of comparison, the cell number in MCAK knockdown or MCAK^-/-^ was normalized to the control.

For wound healing assays, 70 µl of cells at 5-7×10^5^ cells/ml were seeded into a Culture-Insert 2 Well (ibidi) and grown to confluency. After 24 h, the insert was removed, and images of the wound gap were acquired every 2 h over 12 h using a 20x objective (0.60 NA, Plan Apo) on the EVOS microscope. To quantify wound closure, the area (A) of the wound was measured manually using the measure function in Fiji. To compare between experiments, the wound recovery progress (%) was normalized as ΔA/A_0_ for each time point. To calculate the velocity of the wound front, the width of the wound was calculated by dividing the wound area with the height of the wound (image height). The wound width as a function of time was fit with linear regression. The wound front velocity was determined as the slope of the linear fit divided by two.

For the random migration assay, at 30 h post-RNAi transfection, cells were plated at ∼ 2×10^4^/ml on poly-L-lysine coated MatTek dishes and allowed to adhere for 18-24 h. The culture medium was supplemented with 20 mM Hepes (pH 7.2) before imaging. Time-lapse differential interference contrast (DIC) images were obtained every 5 min for 4 h using a 20X objective (0.75 NA, PlanApo) on a DeltaVision microscope (GE Healthcare) equipped with a CoolSNAP HQCCD camera (Photometrics) and controlled by Softworx software (Applied Precision/ GE Healthcare) at 37°C in a humidified chamber. Only individual cells that did not go through mitosis or touch other cells were used for analysis. Cell migration tracks were quantified using a custom semi-automatic MATLAB code. The center of the nucleus of each individual cell over time was manually input, and the x, y coordinates of the nucleus were saved. The migration displacement was defined as the distance between the first (t_1_) and last time point (t_e_), and was calculated as the Euclidian distance 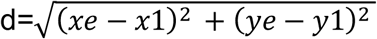. The migration velocity of individual tracked cells was determined as the total distance traveled divided by the total time as d/(t_e_-t_1_). The migration path (p) of a cell was calculated as the sum of distance travelled between each frame, and the directional persistence was calculated as the displacement divided by the migration path d/f (Theisen et al., 2012).

### Cell morphological analysis and centrosome reorientation assay

For cell morphology analysis, at 30 h post-RNAi transfection, cells were plated at ∼2×10^4^ cells/ml on poly-L-lysine coated coverslips and allowed to adhere for 20 h before being processed for immunofluorescence or before being transfected with GFP or CA-Rac1-GFP plasmids. For analysis, cells were segmented using a fluorescence intensity cutoff based on phalloidin-rhodamine or GFP intensity followed by a size cutoff as described above in the analysis of the transwell assay. Cell area and eccentricity (the distance between the foci of the cell/major axis length) of the segmented cells were determined from the MATLAB function regionprops.

For the centrosome reorientation assay, 1.35 ×10^6^ RPE-1 cells were plated in a six-well plate containing poly-L-lysine coated coverslips and were grown for 24 h for full attachment. Cells were transfected for knockdown as described above. At 48 h post-RNAi when cells were grown to full confluency, a scratch wound was introduced to the cells with a 20 µl micropipette tip. Cells were allowed to migrate for 2 h before being processed for immunofluorescence as described above. Single-plane images were collected with a 40x objective (1.0 NA, PlanApo) attached to a Nikon Eclipse 90i microscope equipped with a CoolSnap HQ CCD camera (Photometrics) and controlled by Metamorph (Molecular Devices). The relative position between the centrosome and nucleus was manually quantified as previously described (Chang et al., 2016).

### Focal adhesion turnover assay and analysis in fixed cells

For analysis of FAs in fixed cells, RPE-1 PG6 cells were plated at 1×10^5^ cells/ml onto ploy-L-lysine coverslips in a 6-well plate 24 h prior to RNAi transfection. At 48 h post RNAi transfection, a P20 Pipetman tip was used to create wound edges in the monolayer. After 2 h, cells were fixed with 4% formaldehyde in PHEM (60 mM Pipes, 25 mM Hepes, 5 mM EGTA, 1 mM MgCl, pH 6.9) with 0.2% glutaraldehyde for 20 min. Glutaraldehyde was quenched for 10 min with freshly prepared 0.5 mg/ml NaBH_4_ in PBS. Coverslips were then stained to visualize MTs as described above. Images of cells along the wound edge were captured with a 60X objective (1.42 NA, PlanApo) attached to a Nikon Eclipse 90i microscope equipped with a CoolSnap HQ CCD camera (Photometrics) and controlled by Metamorph (Molecular Devices). A Z-stack was used to capture planes at 0.2 µm steps across a 5 µm range. Images were deconvolved using AutoQuant software (Media Cybernetics).

For analysis of FAs in live cells, at 32 h post RNAi transfection, 100 µl RPE-1 PG6 cells at 5×10^5^/ml were seeded into a Culture-Insert 2 Well placed on 35-mm glass bottom dishes coated with 5 μg/ml fibronectin. Cells were allowed to adhere and grown to confluency for 18-20 h. Hepes (pH 7.2) was added to a final concentration of 20 mM before imaging, and 10–20 image fields were taken along the wound edge every 3 min for 2 h with a 60X objective (1.42 NA, Plan Apo) and captured on a Coolsnap HQ2 camera using a DeltaVision Personal DV microscope.

Analysis of FA turnover was performed as previously described (Kenific et al., 2016; Stehbens and Wittmann, 2014). Only FAs that undergo full turnover from appearance to disappearance during the imaging window were analyzed. Two to five FAs were randomly chosen per cell. To track FAs, ROIs were manually drawn around individual FAs using the Bezier ROI tool in NIS-Elements software (Nikon). The ROI was redrawn if the FA changed size or location over time. The fluorescence intensity of each FA in the time stack was generated using the “Time Measurement” tool. Background was locally corrected by subtracting the fluorescence intensity of a duplicated ROI adjacent to the FA. To smooth the fluorescence intensity curve, a three-frame running average was calculated and plotted as a function of time. The curve was used to calculate the FA assembly rate constant, the disassembly rate constant, and the lifetime by curve fitting using the ‘Solver’ add-in in Excel (Microsoft). Assembly was the first phase of the fluorescence intensity curve when intensity continuously increases and was fit with a logistic function. Disassembly was the latter portion of the intensity plot when fluorescence intensity was decreasing and was fit with an exponential decay function. Some MCAK knockdown cells showed a pause or a second disassembly phase before full disassembly occurred. In these cases, the second disassembly curve was used for fitting to determine the disassembly rate.

### FLIM

For fluorescence lifetime imaging microscopy (FLIM), RPE1 cells were plated at 1.25×10^4^ cells/ml onto poly-L-lysine coated MatTek dishes 24 h before infection and then imaged at 48 h post-infection. An hour prior to imaging, a P20 Pipetmen tip was used to create wound edges within the cell monolayer.

FLIM was performed on a Leica SP8 laser scanning confocal microscope controlled by LAS software (Leica) and equipped with a 40X 1.2 NA HCX PL APO CS water objective (Leica) and an attached PicoQuant multidimensional time-correlated single photon counting system. MatTek dishes containing lentiviral infected cells were placed in a stage-top humidified incubator maintained at 37°C and 5% CO_2_ (Tokai Hit). For confocal imaging, 256 × 256 pixel images were acquired at 400 MHz with a zoom factor of 2, and mEmerald was excited using a white light laser (WLL) at 70% power and the 488 nm laser line at 40 MHz, 70% intensity, and 250% gain. For FLIM imaging, 256 × 256 pixel images were acquired at 200 MHz, and mEmerald was excited by a 488 nm pulsed WLL line at 20 MHz, 70% intensity, and 250% gain until the signal was > 1000 photon counts per pixel. The lifetimes per pixel were determined using SymPhoTime 64 software (PicoQuant) in which an ROI around each cell was manually made by following the contour of the cell, the cell ROI thresholded to 50 counts, the decay data binned 2×2 pixels, and fit to a tri-exponential tail fit decay model that gave a χ^2^ close to 1 (Ems-McClung et al., 2020). The average amplitude-adjusted lifetimes, τ_av/amp_, were then calculated per pixel by fixing background and individual lifetimes determined by the fitting. To calculate the mean lifetime at the leading edge and trailing edge for each cell, the confocal and FLIM images were overlaid, and three 4×4 pixel ROIs were drawn at the leading edge of the cell and three at the rear of the cell body using the confocal image as a positional reference. The mean lifetime for the leading edge or rear of the cell were calculated from each of the three ROIs from the FLIM image.

### Image processing and statistical analysis

Images in each experiment were taken with the same imaging settings, and analyses were done on unprocessed images. Images were processed in Fiji and assembled in Illustrator (Adobe). Data was plotted in MATLAB, Prism 7, or Prism 8 (GraphPad Software), and graphs assembled in Illustrator. Statistical analyses were performed using Prism 7 or Prism 8. To determine if the data came from a normal distribution, a D’Agostino-Pearson omnibus normality test was performed, and a p < 0.05 was considered a non-normal distribution. A normal distribution was assumed when the sample size was small. For comparing two data sets, a two-tailed Student’s *t*-test was used for data distributed normally, and a two-tailed Mann-Whitney U-test was used for non-normal data. For comparing two groups with normal distribution and unequal variances, Welch’s *t*-test was used. For multiple comparisons, an ordinary one-way analysis of variance (ANOVA) followed by Tukey post-hoc test was used for both normal and non-normal distributions. A p < 0.05 was considered significant for all tests.

## Acknowledgements

This work was supported by NIH R35GM122482 to CEW. We thank Anne Straube for Paxillin-GFP expressing RPE-1 cells and Yeajin Kim and Weini Wang for help in optimizing centrosome orientation experiments. We also thank Jim Powers for help with imaging and Sid Shaw for thoughtful discussions of the data and analysis. The IU-LMIC is supported in part by the Office of the Vice Provost for Research.

## Notes

### Competing Interest Statement

The authors have declared no competing interest.

