## Supplemental Figure for "Spatial Regulation of MCAK Promotes Cell Polarization and Focal Adhesion Turnover to Drive Robust Cell Migration"

Supplemental Material

**Figure S1. Knockdown of MCAK alters focal adhesions.** Representative images of control or MCAK knockdown cells expressing paxillin-GFP in which cells were wounded and then allowed to migrate for 2 h. Cells were then stained to visualize MTs (red) and DNA (blue). Images are scaled equivalently. The arrows point in the direction of cell movement toward the wound. Scale bar, 10 µm. Data related to Figure 4.

**Video 1. Representative movies of cells during random migration.** Cells from either control or MCAK knockdown were plated on coverslips, allowed to adhere and then imaged at 5 min intervals for 4 h to examine cell behavior. Movies are recorded at 10 fps. Data related to Figure 2.

**Video 2. Representative movies of focal adhesion dynamics.** Cells from either control or MCAK knockdown were plated in culture inserts in 35 mm glass bottom dishes, allowed to adhere, the insert was removed, and then the cells were imaged along the wound edge at 3 min intervals for 2 h to FA dynamics. Movies are recorded at 10 fps. Data related to Figure 4.

Figure S1

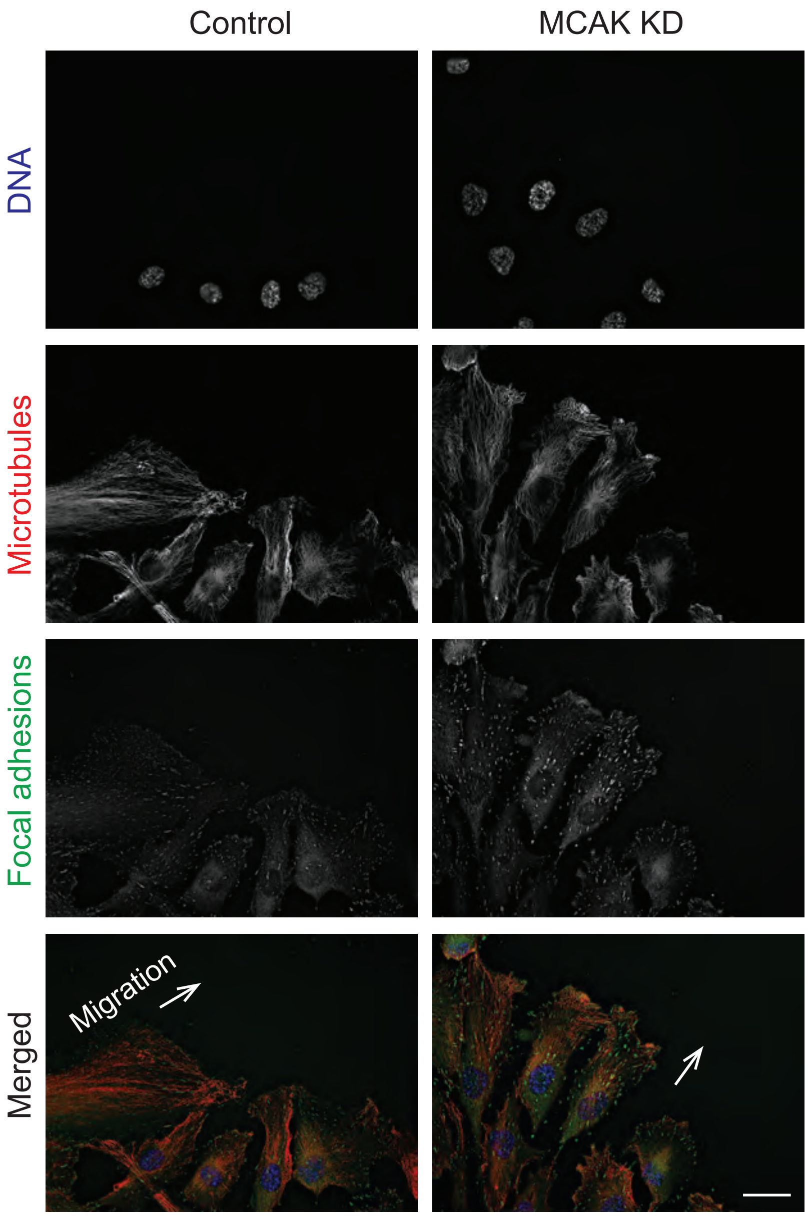

Control
MCAK KD
DNA
Microtubules
Focal adhesions
Merged
Migration
